# Comprehensive genomic analysis of *Klebsiella pneumoniae* and its temperate N-15-like phage: From isolation to functional annotation

**DOI:** 10.1101/2025.03.05.641587

**Authors:** Reham Yahya, Aljawharah Albaqami, Amal Alzahrani, Suha Althubiti, Moayad Alhariri, Eisa T Alrashidi, Nada Alhazmi, Mohammed A. Al-Matary, Najwa Alharbi

## Abstract

A temperate N-15-like phage and an extensively drug-resistant (XDR) Klebsiella pneumoniae strain were studied in this research. The former was found in hospital wastewater, while the latter was retrieved from the sputum of an intensive care unit patient. The bacteria showed strong resistance to several antibiotics, including penicillin (≥16 μg/mL), ceftriaxone (≥32 μg/mL), and meropenem (≥8 μg/mL), which was caused by SHV-11 beta-lactamase, NDM-1 carbapenemase, and porin mutations (OmpK37, MdtQ). Yersiniabactin, enterobactin, and E. coli common pilus (ECP) genes were also present in the genome; these genes are essential for the acquisition of iron, adhesion, and immune evasion, among other virulence factors. kappa testing categorized the strain as K64 and O2a types. The presence of colicin genes, IncHI1B_1_pNDM-MAR and IncFIB replicons, and other plasmids in this strain demonstrate its ability to spread antibiotic resistance and facilitate horizontal gene transfer. Adding to its genetic variety and adaptability, the genome included CRISPR-Cas systems and eleven prophage regions.

The 172,025 bp linear genome and 46.3% GC content of the N-15-like phage showed strong genomic similarities to phages of the Sugarlandvirus genus, especially those that infect *K. pneumoniae.* There were structural proteins (11.8 percent of ORFs), DNA replication and repair enzymes (9.3 percent of ORFs), and a toxin-antitoxin system (0.4 percent of ORFs) encoded by the phage genome. A protelomerase and ParA/B partitioning proteins indicate that the phage is replicating and maintaining itself in a manner similar to the N15 phage, which is renowned for maintaining a linear plasmid prophage throughout lysogeny. Lysogeny and horizontal gene transfer are two mechanisms by which phages may influence bacterial evolution.

Learning about the phage’s role in bacterial evolution, host-phage relationships, and horizontal gene transfer is a great benefit. Understanding the dynamics of antibiotic resistance and pathogen development requires knowledge of phages like this one, which are known for their temperate nature and their function in altering bacterial virulence and resistance profiles. The regulatory and structural proteins of the phage also provide a model for research into the biology of temperate phages and their effects on microbial communities. The importance of temperate phages in bacterial genomes and their function in the larger framework of microbial ecology and evolution is emphasized in this research.

**Author Summary:** Antibiotic-resistant bacteria represent a significant global health threat, and comprehending their interactions with bacteriophages is essential for formulating novel antimicrobial tactics and elucidating the molecular development of bacteria. This work examined an extensively drug-resistant (XDR) strain of *Klebsiella pneumoniae* isolated from an ICU patient and a temperate N-15-like phage identified in hospital effluent. The bacterial strain exhibited resistance to multiple antibiotics owing to an array of resistance genes, plasmids, porin mutations, and virulence characteristics that facilitated its survival and pathogenicity. The genomic investigation of the phage elucidated its structural organization, replication mechanisms, and possible contribution to bacterial development through lysogeny and horizontal gene transfer. Our findings underscore the intricate host-phage interactions that affect antibiotic resistance and pathogenicity in K. pneumoniae, offering significant insights into microbial evolution and the prospective relevance of phages in therapeutic approaches.

## Introduction

The development of extensively drug-resistant (XDR) bacterial pathogens poses a major threat to public health worldwide. Now a major pathogen causing hospital-acquired infections, including urinary tract infections, pneumonia, and bloodstream infections, is *Klebsiella pneumoniae*. *K. pneumoniae*’s repute as a major nosocomial pathogen has been much boosted by its ability to acquire and disseminate antibiotic resistance genes via horizontal gene transfer routes, including plasmids, transposons, and prophages (1). Antibiotic resistance genes (ARGs), especially those that code for beta-lactamases and carbapenemases, have made many first-line antibiotics less effective. This makes treatment plans more difficult and puts more stress on healthcare systems (2). Virulence factors, including siderophores (e.g., yersiniabactin and enterobactin) and adhesion systems (e.g., *E. coli* common pilus), increase the bacterium’s potential to colonize host tissues, avoid immune responses, and start chronic infections (3). Developing sensible plans to treat *K. pneumoniae* depends on an understanding of the genetic pathways leading to antibiotic resistance and virulence. By means of horizontal gene transfer (HGT) and manipulation of bacterial virulence and resistance traits, bacteriophages, also known as phages, much affect bacterial evolution (4). Temperate phages mix with bacterial genomes as prophages and may contain genes that provide their hosts selection benefits, such as antibiotic resistance or virulence factors (5). The N15-like phage is a temperate phage whose genome remains a linear plasmid after lysogeny (6). Protelomerase and partitioning proteins (ParA/B) help the phage to survive among bacterial populations and increase their genetic variation (7). The linear genome of the N15-like phage distinguishes it from most other temperate phages that do not merge into the host chromosome; it survives as a linear plasmid during the lysogenic stage. This replication system guarantees consistent phage genome inheritance and helps bacterial hosts to horizontally transmit genetic material, including ARGs and virulence factors (8). Researching these phages helps one to grasp the dynamics of host-phage interactions and the processes behind phage-mediated bacterial evolution. It codes for a lot of different proteins, such as structural proteins, DNA replication and repair enzymes, and toxin-antitoxin systems that make it easier for the genome to infect, multiply, and stay inside bacterial hosts (9). Protelomerase, which changes the linear phage genome into a circular form during replication, and partitioning proteins, which help the phage genome to be stably segregated during cell division, highlight the several mechanisms the phage uses to preserve its genetic material (7). The features of the N15-like phage make it a useful model for examining the biology of temperate phages and their impact on microbial populations (8). Understanding bacterial evolution, host-phage interactions, and the spread of antibiotic resistance depends on an analysis of temperate phages (4). Understanding the processes of antibiotic resistance and pathogen development requires an awareness of the part phages play in increasing the genetic diversity and adaptability of bacterial pathogens (5). Studying the biology of temperate phages and their consequences on microbial populations is modeled by the structural and regulatory proteins of the phage (8). Temperate phages are good candidates for phage treatment and phage engineering because of their unique qualities, including their ability to integrate into bacterial genomes and their aptitude to carry beneficial genes. Because lytic phages may directly kill bacterial hosts, they have been favored for therapeutic uses. Temperate phages, on the other hand, provide special advantages, including the capacity to remain inside bacterial populations and to transfer genetic material that can compromise bacterial virulence or antibiotic resistance (10). Synthetic biology has recently advanced the engineering of temperate phages to increase their medicinal effectiveness. Designed to particularly target and destroy antibiotic resistance genes or virulence components in bacterial genomes, temperate phages may be made to integrate CRISPR-Cas systems (11). Moreover, temperate phages may be altered to remove integration machinery, therefore changing them into "lysis-only" phages that retain the ability to infect and lyse bacterial hosts without establishing lysogenic connections (12). Engineering projects find a strong foundation in the N15-like phage, distinguished by its unique linear plasmid replication process. By use of partitioning proteins and the protelomerase system, scientists may design creative phage-based treatments that combine the lytic characteristics of traditional therapeutic phages with the stability of temperate phages. Research of temperate phages like N15 provides valuable information for the creation of synthetic phages meant to especially target bacterial infections, hence minimizing off-target consequences. As phage treatment develops, temperate phages should play a major part in the evolution of next-generation antimicrobial approaches (13). The present investigation examines an N15-like phage identified in hospital wastewater and an extensively drug-resistant *K. pneumoniae* strain obtained from a clinical environment. The genetic and functional traits of the bacterial strain and the phage investigated in this study to learn more about how bacteria and phages interact and what that means for pathogen evolution and antibiotic resistance. This study underscores the significance of temperate phages in bacterial genomes and their contributions to microbial ecology and evolution, thereby indicating the necessity for ongoing research into phage-bacteria interactions to combat antibiotic-resistant diseases.

## Results

### Genome assembly and strain phylogenetic analysis

The total genome assembly of the strain was 5,814,374 bp with a GC content of 56.84% was deposited in GenBank database under bioproject number PRJNA1217456, biosample number SAMN46489325 and assembly accession number GCA_047532915.1. The assembly was of high quality, with 0.00 N’s per 100 kbp, indicating no ambiguous bases in the 175 contigs. These metrics collectively demonstrate a robust and continuous genome assembly, suitable for downstream analysis. MLST analysis revealed that the strain belongs to ST147 high risk clone according to Pasteur database.

The phylogenetic investigation (Fig 1) indicates a close evolutionary link between *Klebsiella pneumoniae* kpn_R01 and the clinical isolate CP120872 from China. Clinical isolates CP148957 from Japan and CP119564 from the United Kingdom further cluster with this strain, suggesting potential global transmission. The extended family tree includes CP063927, a Chinese environmental isolation; and CP083775, an Australian clinical isolate. The proximity of environmental and clinical isolates indicates that environmental reservoirs contributed to the transmission of the lineage. The identification of analogous strains in China, Japan, the United Kingdom, and Australia contributes to the evidence of potential global dissemination. This may result from infections acquired during hospital care or while patients are abroad.

**Fig 1.**
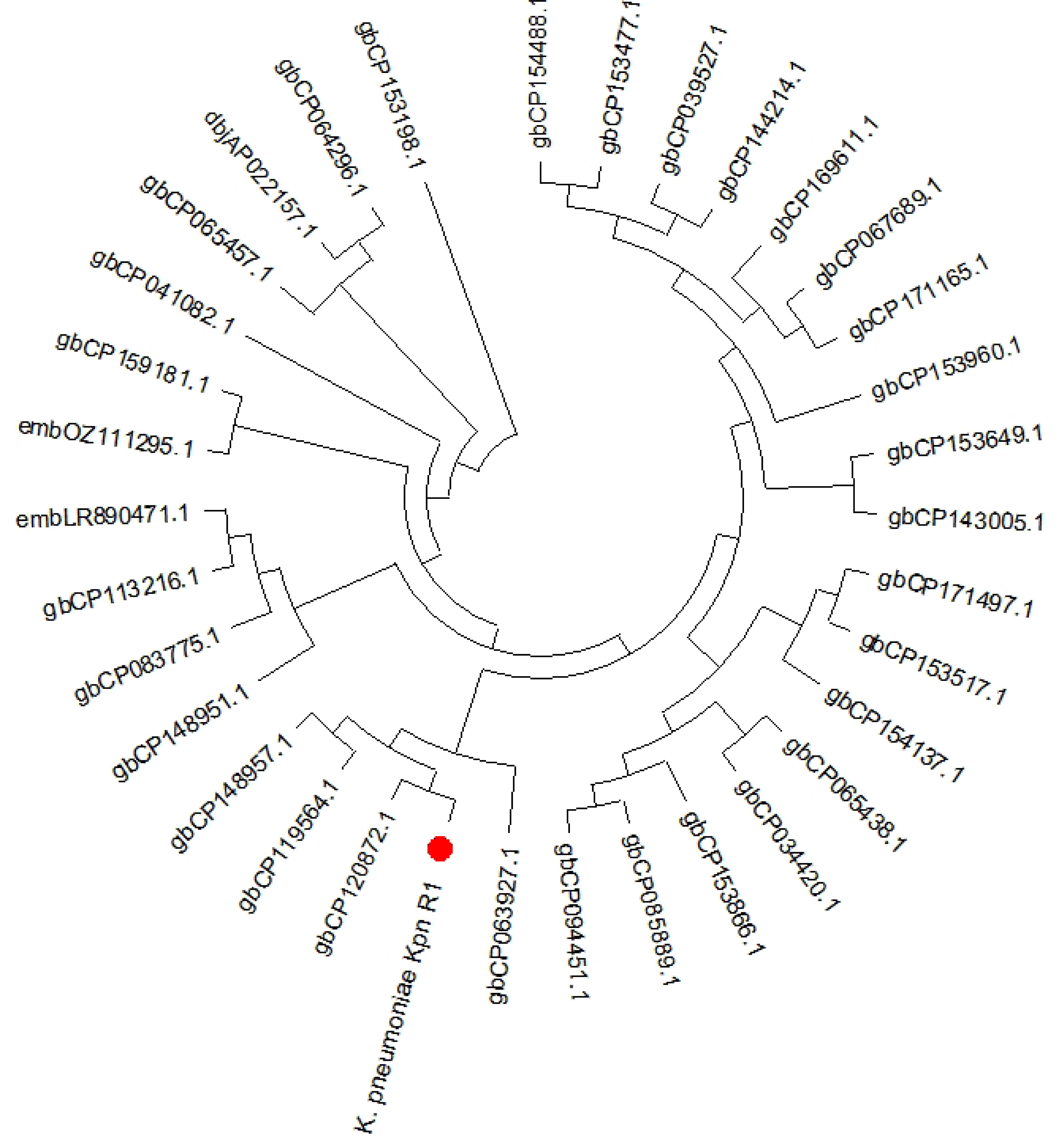
Circular phylogenetic tree depicting the evolutionary relationships of *Klebsiella pneumoniae* Kpn_R01 and its closest 34 strains. The Maximum Likelihood (ML) tree was constructed to illustrate the phylogenetic placement of *K. pneumoniae* Kpn_R01 (highlighted with a red dot) relative to closely related strains. The clustering pattern suggests potential global dissemination, with strains from various geographic locations. The close association between clinical and environmental isolates highlights the role of environmental reservoirs in bacterial transmission.

**Fig 2.**
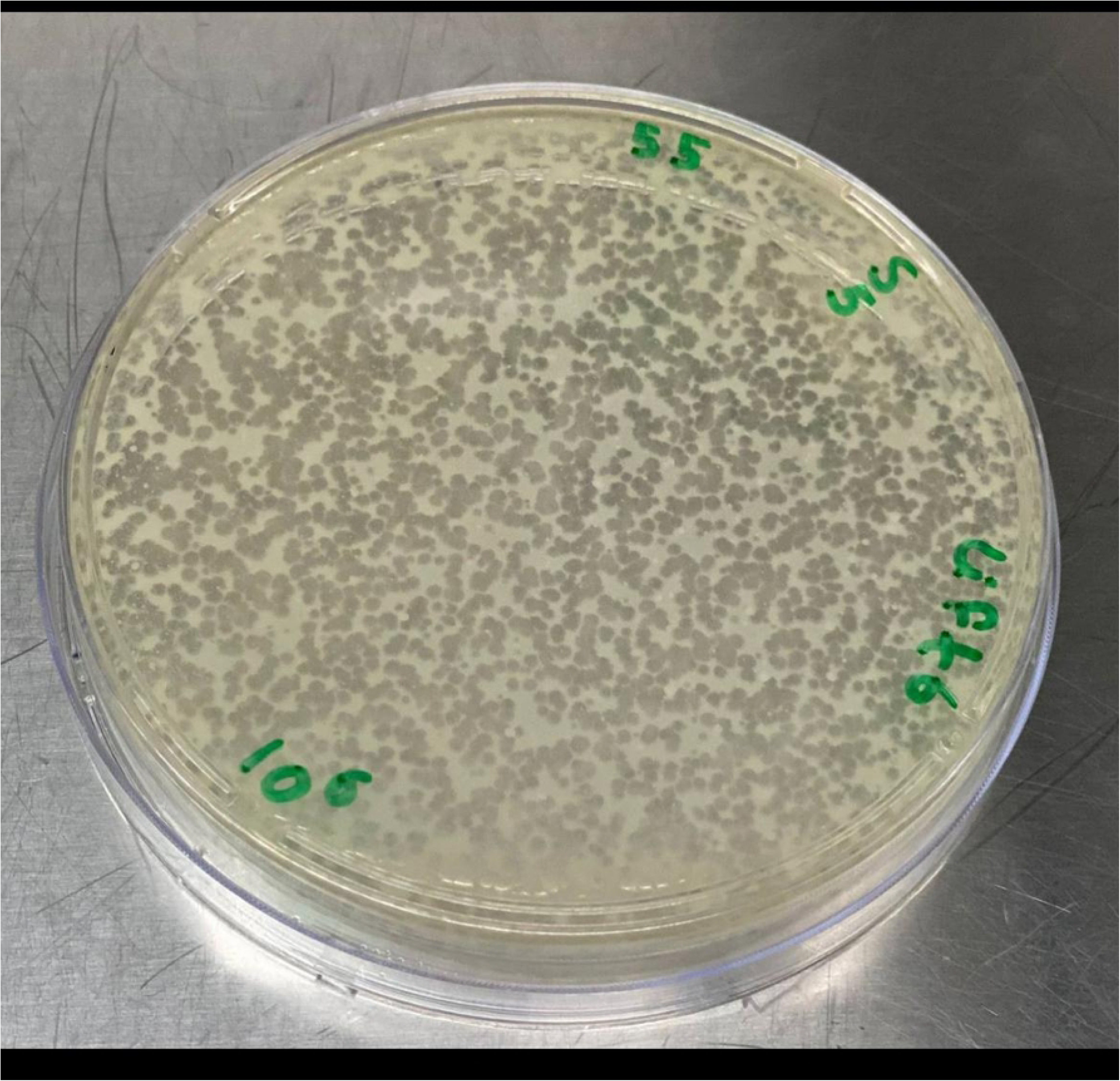
**Isolation and characterization of Klebsiella phage Kpn_R1.** (A) Plaque morphology of Klebsiella phage Kpn_R1 on a bacterial lawn, showing distinct, clear plaques. (B) Transmission electron microscopy (TEM) image of Klebsiella phage Kpn_R1, revealing a polyhedral head (53.7 nm in diameter) and a wavy tail (178.3 nm in length). Based on morphology, the phage is classified within the *Demerecviridae* family according to the International Committee on Taxonomy of Viruses (ICTV). Scale bar: 200 nm.

### Antimicrobial susceptibility and resistome

The strain’s XDR trait can be seen in the antibiotic-resistant genes found in the composite genome and analyzed using the CARD database. The genes possess distinct resistance mechanisms that counteract various drug types. As shown in Table 1, SHV-11 beta-lactamase, NDM-1 carbapenemase, and mutations in porines (OmpK37, MdtQ) mediate the strain resistance to beta-lactam antibiotics, resulting in minimum inhibitory concentrations (MICs) of penicillin (≥16 μg/ml), ceftriaxone (≥32 μg/ml), and meropenem (≥8 μg/ml). Aztreonam resistance (≥32 μg/ml) is similarly attributed to NDM-1 and low permeability. Minimum inhibitory concentrations (MICs) of ciprofloxacin (≥4 μg/ml) and levofloxacin (≥8 μg/ml) are primarily due to fluoroquinolones is augmented by efflux pumps (oqxA, acrA, marA) and target mutations (rsmA). Enhanced aminoglycoside resistance is ascribed to 16S rRNA methyltransferase (armA) and aminoglycoside-modifying enzymes (aadA2), leading to minimum inhibitory concentrations (MICs) of ≥16 μg/mL for gentamicin and ≥64 μg/mL for amikacin. Resistance to tetracycline (≥16 μg/mL) and tigecycline (≥2 μg/mL) is facilitated by efflux pumps (tet(A) and KpnF). Efflux pumps (KpnF, KpnG, KpnH) and enzymatic inactivation (msrE, mphE) promot resistance to Macrolide, resulting in minimum inhibitory concentrations (MICs) of ≥8 μg/mL for erythromycin and ≥16 μg/mL for azithromycin. Resistance to colistin and polymyxin B (≥4 μg/mL) is associated with phosphoethanolamine transferase (ArnT, eptB) and reduced permeability (OmpA). Increased resistance to sulfamethoxazole (≥256 μg/mL) and trimethoprim (≥16 μg/mL) is ascribed to target replacement (sul1, dfrA12). Chloramphenicol resistance (≥32 μg/mL) is mediated by efflux pumps (acrA, rsmA), while fosfomycin resistance (≥64 μg/mL) arises from enzymatic inactivation (fosA5). Moreover, resistance to quaternary ammonium compounds (≥50 μg/mL) is facilitated by efflux mechanisms (qacEdelta1, leuO). The extensive resistance profile of this strain highlights the challenges in managing infections caused by XDR *K. pneumoniae* and underscores the need for alternative treatment strategies.

**Table 1.** The Minimum Inhibitory Concentrations (MICs) for various antimicrobials and the resistance mechanisms identified in the strain.

| Antimicrobial Class | Antimicrobial Agent | MIC ($\mu\text{g/mL}$) | Gene | Resistance Mechanism | Notes |
| --- | --- | --- | --- | --- | --- |
| Beta-Lactams | Penicillin | $\geq 16$ | SHV-11<br>NDM-1<br>OmpK37, MdtQ | beta-lactamase<br>carbapenemase<br>reduced permeability | High resistance due to beta-lactamase and porin mutations. |
| Cephalosporin | ceftriaxone | $\geq 32$ | SHV-11<br>NDM-1<br>OmpK37, MdtQ | reduced permeability | Resistance to broad-spectrum cephalosporins. |
| Carbapenem | meropenem | $\geq 8$ | NDM-1<br>OmpK37, MdtQ | reduced permeability | Carbapenem resistance due to carbapenemase and porin loss. |
| Monobactam | aztreonam | $\geq 32$ | NDM-1<br>MdtQ | reduced permeability | Resistance due to carbapenemase activity |
| Fluoroquinolones | Ciprofloxacin | $\geq 4$ | oqxA, acrA, marA, rsmA<br>marA | Efflux pumps<br>reduced permeability | High resistance due to efflux and target mutations |
| | Levofloxacin | $\geq 8$ | oqxA, acrA, marA, rsmA | Efflux pumps | Similar to ciprofloxacin resistance. |
| Aminoglycosides | Gentamicin | $\geq 16$ | armA<br>aadA2 | 16S rRNA methyltransferase<br>aminoglycoside-modifying enzyme | High-level resistance due to target modification and enzymatic inactivation |
| | Amikacin | $\geq 64$ | armA | 16S rRNA methyltransferase | Resistance due to target modification |
| Tetracyclines | Tetracycline | $\geq 16$ | tet(A)<br>*K. pneumoniae*<br>KpnF | efflux pump<br>efflux pump | Efflux-mediated resistance |
| | Tigecycline | $\geq 2$ | tet(A)<br>*K. pneumoniae*<br>KpnF | efflux pump<br>efflux pump | Tigecycline resistance is rare but possible due to efflux mechanisms |
| Macrolides | Erythromycin | $\geq 8$ | msrE<br>mphE<br>KpnF, KpnG, KpnH | ABC-F protein<br>macrolide phosphotransferase<br>efflux pumps | Resistance due to efflux and enzymatic inactivation. |
| | Azithromycin | $\geq 16$ | msrE<br>mphE<br>KpnF, KpnG, KpnH | efflux pumps | Similar to erythromycin resistance. |
| Peptide Antibiotics | Colistin (Polymyxin E) | $\geq 4$ | ArnT<br>eptB<br>OmpA | phosphoethanolamine transferase<br>reduced permeability | Resistance due to target modification and reduced permeability |
| | Polymyxin B | $\geq 4$ | | | |
| Sulfonamides | Sulfamethoxazole | $\geq 256$ | sul1 | sulfonamide-resistant dihydropteroate synthase | High resistance due to target replacement |
| Diaminopyrimidines | Trimethoprim | $\geq 16$ | dfrA12 | trimethoprim-resistant dihydrofolate reductase | Resistance due to target replacement |
| Phenicol | Chloramphenicol | $\geq 32$ | acrA, rsmA | Efflux pumps | Resistance due to efflux mechanisms |
| bleomycin |  |  | BRP(MBL) |  | Resistance due to target alteration and enzymatic inactivation. |
| Rifamycins | Rifampin | $\geq 8$ | KpnF, KpnE, marA | Efflux pumps | Resistance due to efflux mechanisms |
| Fosfomycin | Fosfomycin | $\geq 64$ | fosA5 | fosfomycin thiol transferase | Resistance due to enzymatic inactivation |
| Disinfectants/Antiseptics | Quaternary ammonium compounds | $\geq 50$ | qacEdelta1<br>leuO | efflux pump | Resistance due to efflux mechanisms |

### Genes linked to virulence

Table 2 shows that *K. pneumoniae* Kpn_R01 strain harbours a number of important virulence factors that were identified via the examination of virulence-associated genes. These factors included genes linked to the production of yersiniabactin, enterobactin, and the *E. coli* common pilus (ECP). High coverage (98.31% to 100%) and identity (85.28% to 90.07%) of the ECP gene cluster (ecpR, ecpA, ecpB, ecpC, ecpD, ecpE) indicates that the strain may manufacture E. coli common pili, which are important in adhesion and biofilm formation. This strain has the ability to produce yersiniabactin, a siderophore that aids in iron acquisition and virulence; the yersiniabactin biosynthetic gene cluster (fyuA, ybtE, ybtT, ybtU, irp1, irp2, ybtA, ybtP, ybtQ, ybtX, ybtS) was also detected with 100% coverage and 97.62% to 99.95% identity. The strain has the ability to produce enterobactin, a siderophore that plays a crucial role in iron absorption. Genes involved in its production were identified (fepC, fepG, entB, entA), the coverage and identity were (88.72% to 99.33%) and (80.00% to 82.65%), respectively. In addition to being discovered with 100% coverage and 83.75% identity, the ompA gene codes for the outer membrane protein A; hence, this strain may be able to elude the immune system and cause diseases. Among these outcomes are the strain’s pathogenicity-critical capabilities— its ability to adhere, absorb iron, and evade the immune system.

**Table 2.**
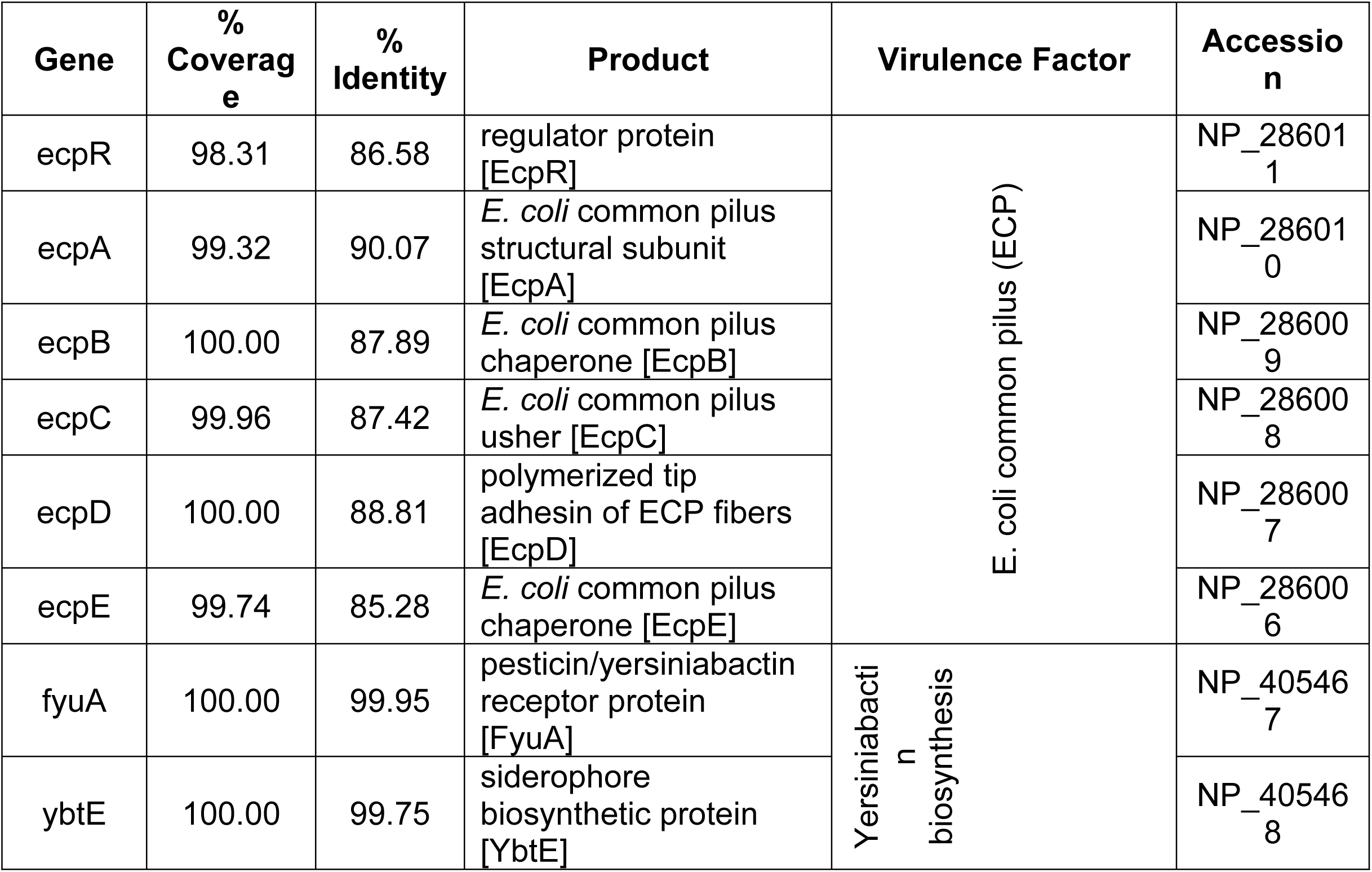
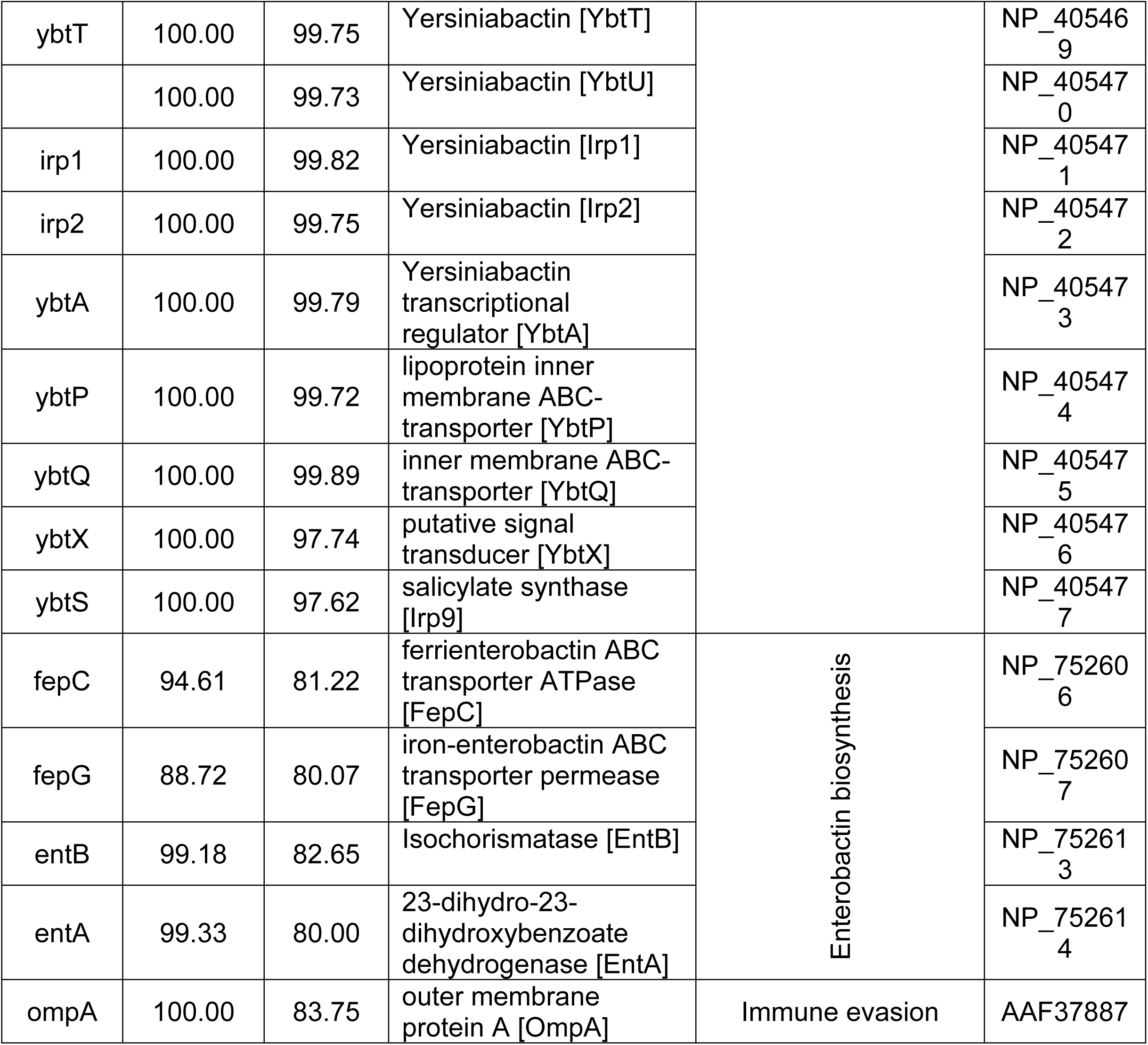
Virulence-associated genes identified in the strain.

### K- and O serotypes

According to the results of Kaptive analysis shown in Table 3, there are two distinct loci that correspond to the K64 and O2a serotypes, respectively: KL64 and O1/O2v1. With 100% identity and 90.10% coverage, indicating a partial match, the KL64 locus was determined to be typeable. Out of the 24 anticipated genes in the KL64 locus, 23 were identified (95.83%), with one gene (KL64_12_wcoT) absent. Numerous genes within the KL64 locus were either partly shortened or incomplete, including wzx, wcoV, wzy, wcoU, and wcsF, potentially impacting the functioning of the capsular polysaccharide production pathway. Furthermore, three other genes outside the locus were found, namely rfaG, gmd, and HG290, which are linked to other serotypes (KL40, KL150, and KL4).

**Table 3.** The key findings from the assembly analysis O-antigen and capsular type.

| Category | O1/O2v1 Locus | K64 Locus |
| --- | --- | --- |
| Best Match Locus | O1/O2v1 | KL64 |
| Best Match Type | O2a | K64 |
| Match Confidence | Typeable | Typeable |
| Identity | 99.03% | 100.00% |
| Coverage | 100.00% | 90.10% |
| Expected Genes in Locus | 7 / 7 (100.00%) | 23 / 24 (95.83%) |
| Expected Gene Details | wzm (100%, 100%);<br>wzt (100%, 100%);<br>wbbM (99.21%, 100%);<br>glf (97.92%, 100%);<br>wbbN (97.31%, 100%);<br>wbbO (99.20%, 100%);<br>kfoC (99.54%, 100%) | galF (100%, 100%);<br>cpsACP (100%, 100%);<br>wzi (100%, 100%);<br>wza (100%, 100%);<br>wzb (100%, 100%);<br>wzc (100%, 100%);<br>wzx (100%, 81.85%, partial);<br>wcoV (100%, 23.45%, partial);<br>wzy (100%, 79.16%, partial);<br>wcoU (100%, 63.69%, partial);<br>wcsF (100%, 39.57%, partial);<br>wcuK (100%, 100%);<br>wbaZ (100%, 100%);<br>wcaJ (100%, 100%);<br>gnd (100%, 100%);<br>manC (100%, 100%);<br>manB (100%, 100%);<br>rmlB (100%, 100%);<br>rmlA (100%, 100%); |
|  |  | rmID (100%, 100%);<br>rmIC (100%, 100%);<br>ugd (100%, 100%) |
| Missing Expected Genes | None | KL64_12_wcoT |
| Other Genes in Locus | None | None |
| Expected Genes Outside Locus | 0 / 7 (0.00%) | 0 / 24 (0.00%) |
| Other Genes Outside Locus | 6 | 3 |
| Other Gene Details | manC (100%, 100%);<br>manB (99.77%, 95.65%);<br>rfbB (98.02%, 100%);<br>rfbD (98.65%, 100%);<br>rfbA (97.19%, 98.62%);<br>rfbC (96.69%, 98.37%) | rfaG (98.42%, 99.74%);<br>gmd (98.80%, 100.00%);<br>HG290 (80.19%, 88.25%) |
| Truncated Genes | None | wzx, wcoV, wzy, wcoU, wcsF |
| Extra Genes | None | None |

The O1/O2v1 locus indicated that the strain was accurately categorized as O2a, exhibiting 99.03% identity and 100% coverage. The seven O-locus genes (wzm, wzt, wbbM, glf, wbbN, wbbO, and kfoC) were predicted as present and undamaged with high identity and coverage ranging from 97.31% to 100%. In addition to the locus, six other genes were identified: manC, manB, rfbB, rfbD, rfbA, and rfbC. These genes are often associated with different serotypes, such as O3b, O12, and OL13. These results indicate a highly preserved O2a serotype with possible genetic interchange with other serotypes. The strain exhibits a well-preserved O2a serotype along with supplementary genetic components that may enhance its antigenic variety and indicate a possible genetic exchange with other serotypes.

### Plasmids

Several plasmid replicons and colicin genes were found in the strain’s examination of plasmid-associated genes, suggesting that the plasmid content is varied (Table 4). A plasmid linked to NDM carbapenemase resistance may be present in the IncHI1B_1_pNDM-MAR replicon, which was found with 100% coverage and identity. Also present were IncFIB (K)_1_Kpn3 and IncFIB (pKPHS1) _1_pKPHS1 replicons, which are often associated with *K. pneumoniae,* with 100% coverage and 95.54% and 91.07% identity, respectively. The presence of high coverage (100%) and identity (88.60% to 100%) for colicin genes, such as Col440I_1, Col (BS512)_1, and ColpVC_1, indicates that this strain may be capable of producing colicins, bacteriocins that may suppress competing bacterial strains. These results demonstrate that this strain might be capable of horizontal gene transfer and the spread of antibiotic resistance by use of plasmids.

**Table 4.**
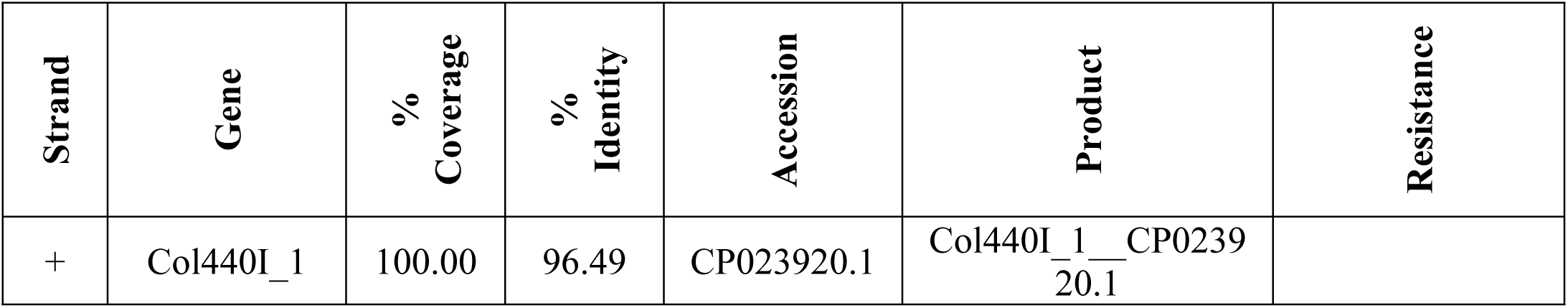
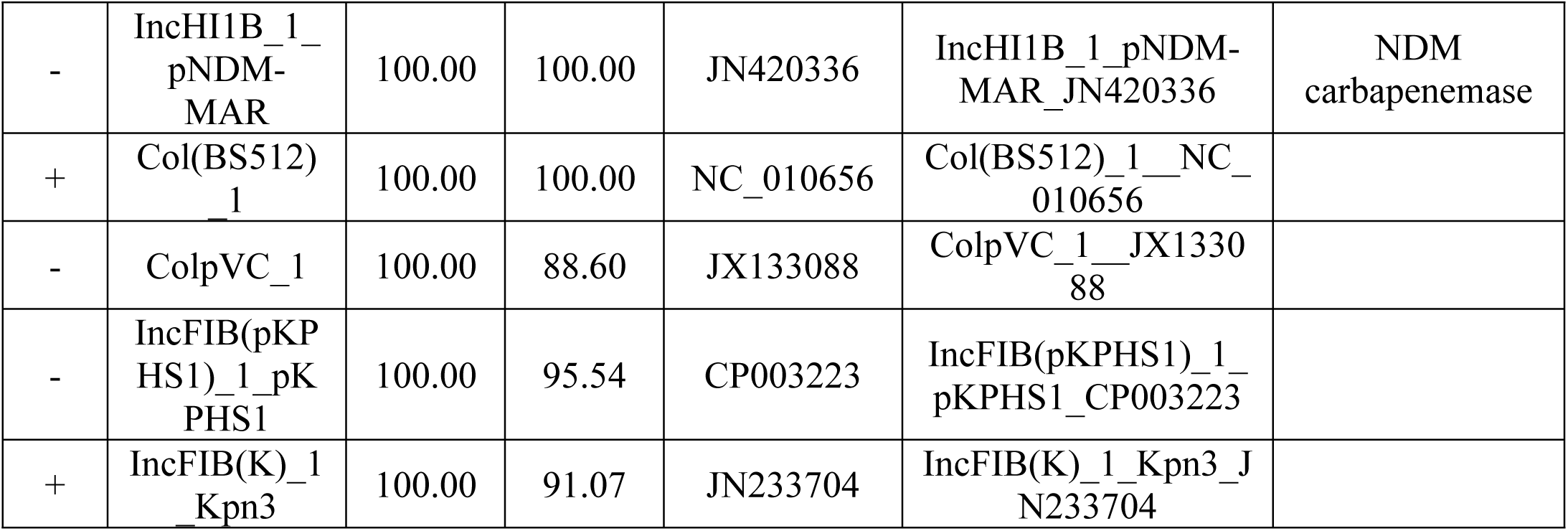
Plasmids Predicted from Contigs of Whole Genome Sequencing (WGS) of *Klebsiella pneumoniae* Kpn_R01 Strain Using PlasmidFinder.

### CRISPR-Cas Systems

The CRISPR-Cas Finder analysis revealed unique CRISPR arrays in two sequences, Seq31 and Seq37. Seq31 consisted of a single CRISPR array (Seq31_1) with a length of 139 bp, featuring 50-bp repeats and 40-bp spacers, demonstrating 98% conservation in both the repeats and spacers. Seq37 exhibited a significantly larger CRISPR array (Seq37_1) of 2,649 bp, consisting of 29-bp repeats and 43 spacers, with a repeat conservation of 75.86% and a spacer conservation of 95.69%. Seq37 was oriented in the forward direction and exhibited an evidence level of 4, indicating a well-defined CRISPR locus. The findings demonstrate that although CRISPR elements are present in the dataset, their prevalence is low, with only a limited number of sequences showing functional or well-conserved CRISPR arrays (S1 Table).

### Prophage Regions and Viral Sequences

VirSorter analysis detected 14 potential prophage sequences in the sample, the majority of which were dsDNA phages, as well as one ssDNA phage (Seq129). The prophage sequences varied in length from 2,300 bp (Seq129) to 49,050 bp (Seq34). The highest confidence values (maximum score = 1.00) were found in Seq34 and Seq43, indicating strong prophage signals. The number of signature genes ranged from 0 to 14, with Seq34 having the most. Multiple sequences, including Seq86 and Seq102, had low confidence scores (0.613 and 0.50, respectively) and lacked hallmark genes, indicating weaker prophage signatures. The discovery of these putative prophages provides information about potential viral components inside the investigated bacterial genomes, which may influence bacterial evolution and resistance mechanisms (S2 Table).

### *Klebsiella* phage Kpn_R1 isolation, morphology, assembly and taxonomy

The phage was isolated from hospital wastewater with distinct plaque morphology (Fig 1A). TEM image examination showed that *Klebsiella* phage Kpn_R1 had polyhedral head with a diameter of 53.7 nm and a wavy 178.3 nm long tail morphology (Fig 1B). This, pursuant to the International Committee on Taxonomy of Viruses taxonomy system, places them in the Demerecviridae family.

### General Genome Characteristics

The phage genome is composed of **172025 bp** base pairs (bp), with a GC content of 46.3%. The genome consists of 211 predicted genes, with an average gene length of **639.47 bp**. The genome type is classified as unknown, and it is of high-quality with a 100.0% genome quality score with 100.0% completeness and 0.0% contamination.

The estimated temperate of the phage is 81%, based on the BACPHLIP database. No antibiotic resistance genes were identified within the genome, indicating that the phage does not harbour any known antibiotic resistance traits, and the phage is confirmed not to be a provirus. Phage genome organization is shown in Fig 3. The phage whole genome sequencing contigs were deposited to GenBank under submission numbers PQ800144.1-PQ800159.1.

**Fig 3.**
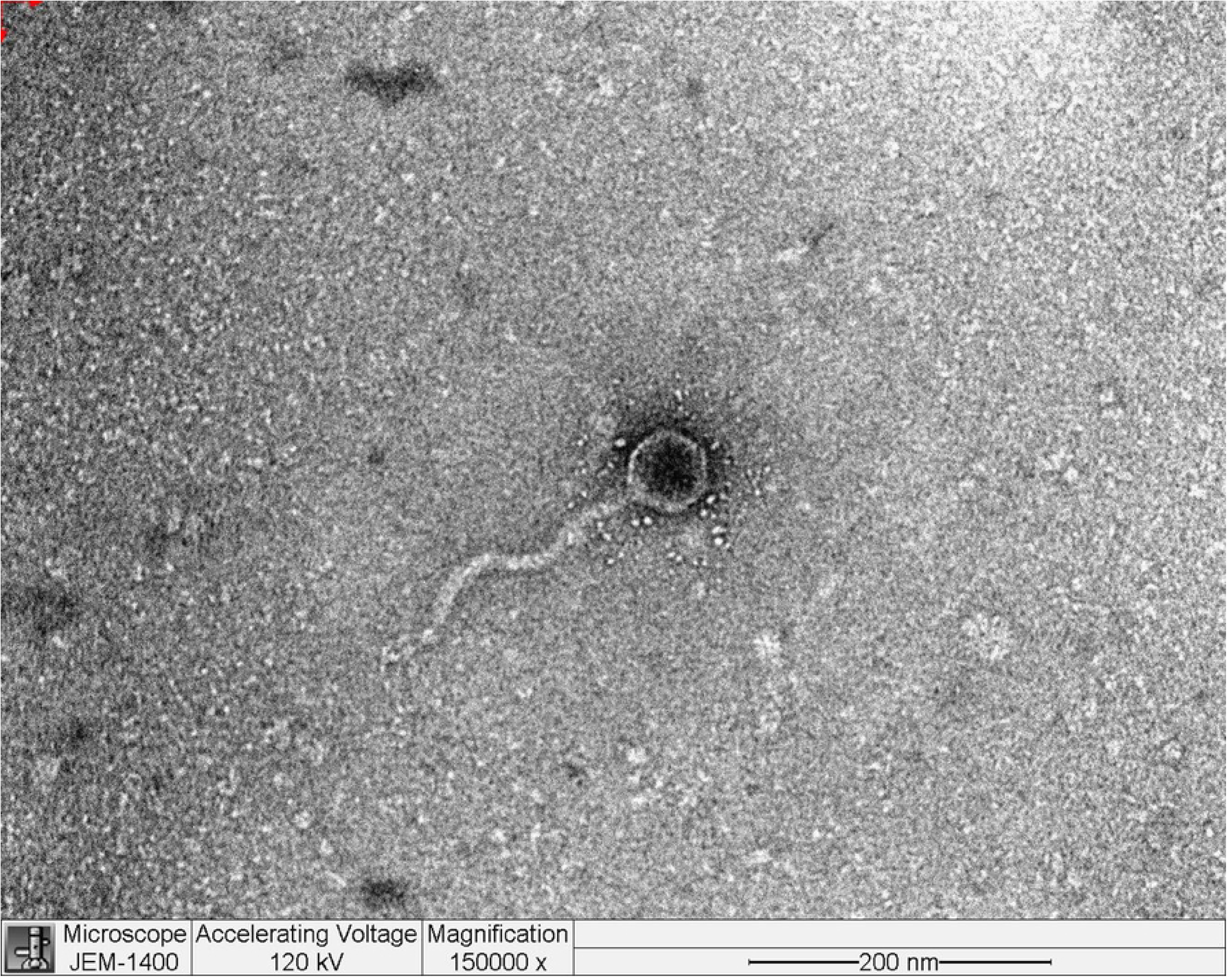
Genome organization of *Klebsiella* phage Kpn_R1. The phage genome is 172,025 base pairs (bp) in length with a GC content of 46.3%. The genome consists of 220 predicted genes, with an average gene length of 639.47 bp. Genes are represented as arrows indicating their orientation. No antibiotic resistance genes were identified, and the phage was confirmed not to be a provirus.

### Predicted Host Range, Evolutionary Relationships and Genomic Similarity of *Klebsiella* Phage Kpn_R1

The predicted hosts of the annotated phage were inferred based on the best-matching phage genomes and their reported hosts. As shown in Table 5, the majority of hits (33 hits) were associated with *Klebsiella*, particularly *K. pneumoniae,* with an average genetic distance of 0.06, an average matching hash score of 215.3 and an average p-value of hits 3.38e-111. This strong association suggests that *K. pneumoniae* is the most likely host for the annotated phage. Other potential hosts include *Yersinia* (2 hits), *Salmonella* (13 hits), *Erwinia* (1 hit), and Campylobacter (1 hit), though these associations are weaker, as indicated by higher genetic distances (0.22–0.23).

**Table 5.**
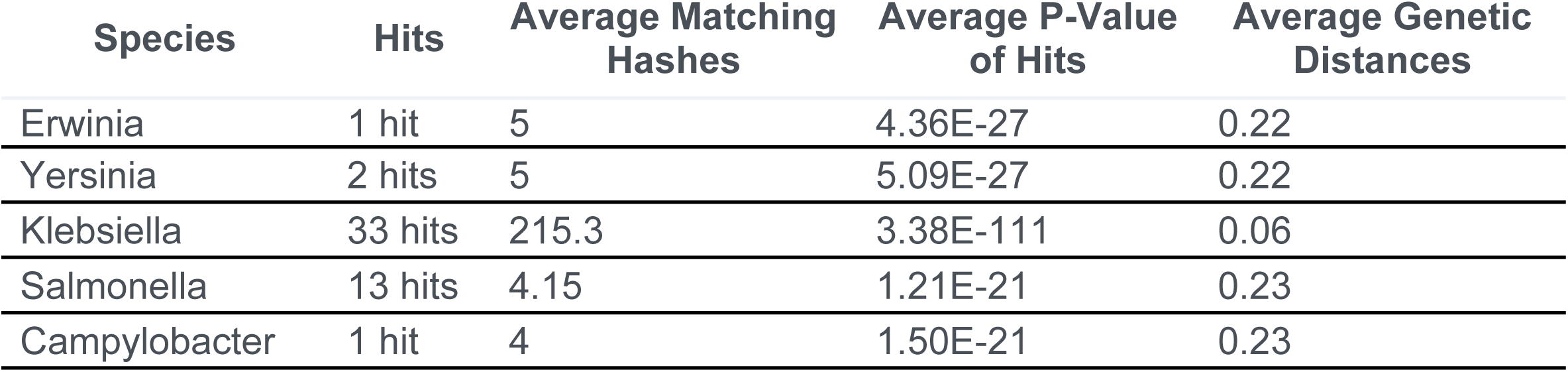
Predicted hosts as inferred through the best-matching phage genomes and their reported hosts.

Table 6 shows that the analysis grouped the closest relatives into several genera, with the highest number of hits belonging to Sugarlandvirus (20 hits) and Epseptimavirus (15 hits). The Sugarlandvirus genus showed the lowest average genetic distance (0.04) and the highest average matching hashes (250.8), indicating a strong genomic similarity. These results suggest that the isolated phage belongs to Caudoviricetes; Demerecviridae; Sugarlandvirus genus.

**Table 6.**
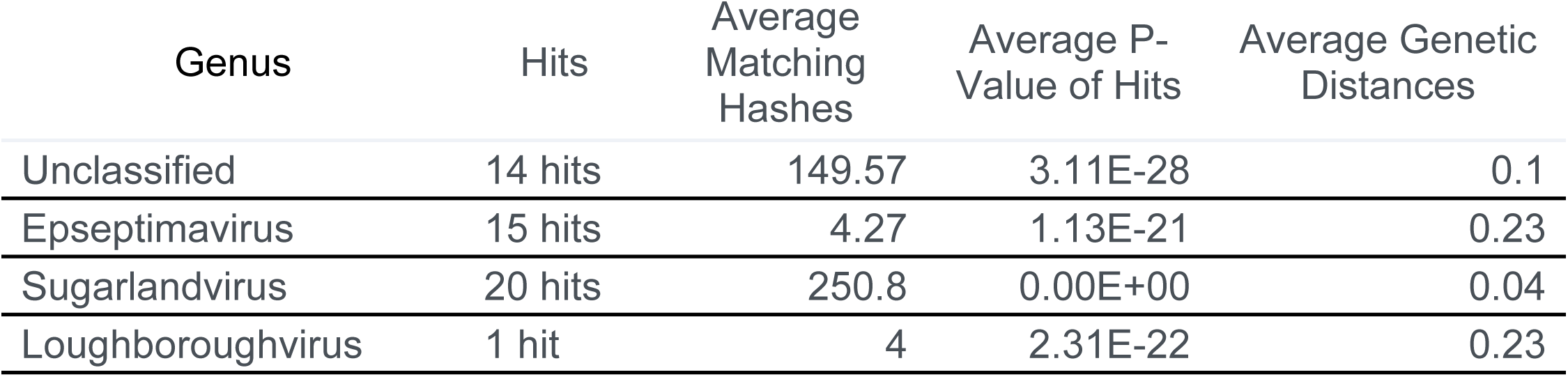
Taxonomic classification and genetic similarity metrics of *Klebsiella* phage Kpn_R1 and its closest relative genera (using the MASH algorithm), All hits with genetic distance < 1 against published phage genomes grouped by genus.

The evolutionary relationships of Klebsiella phage Kpn_R1 were investigated using whole-genome sequencing comparisons (Fig. 4). The circular phylogenetic tree shows different clusters of closely related phages, with Kpn_R1 located next to Klebsiella phage vB_Kpn_IME260 and Klebsiella phage Sugarland. Kpn_R1 shares 92.8% genomic identity across 31.5% of its sequence with vB_Kpn_IME260, as well as 92.3% identity over 30.1% of its genome with Klebsiella phage Sugarland. The study revealed Escherichia phage N15, which had a mean identity of 69.8% across 14% of the genome, indicating a significant evolutionary difference.

**Fig. 4.** Phylogenetic analysis of *Klebsiella* phage Kpn_R1. The evolutionary relationships of Kpn_R1 were analyzed using whole-genome sequence comparisons. The circular phylogenetic tree, constructed using the **maximum likelihood** method, illustrates the closest related phages forming distinct clusters (gray). Kpn_R1 shares 92.8% identity across 31.5% of its genome with *Klebsiella* phage vB_Kpn_IME260 and 92.3% identity across 30.1% of its genome with *Klebsiella* phage Sugarland. The comparison also identified *Escherichia* phage N15 with a lower mean identity (69.8% across 14% of the genome), suggesting evolutionary divergence. Seq1 (marked with red branch) represent *Klebsiella* phage Kpn_R1

The alignment of protein sequences (Fig. 5) further highlights the similarities between Kpn_R1 and its closest relatives. The strongest sequence similarities between Klebsiella phage Kpn_R1 (Seq1), Klebsiella phage vB_Kpn_IME260, and Klebsiella phage Sugarland are depicted by connected syntenic blocks, with the most conserved regions shown in pink. The distinct structural organization of Escherichia phage N15 further reinforces its evolutionary divergence.

**Fig. 5.** Proteomic similarity map of Klebsiella phage Kpn_R1 and related phages. The alignment shows protein sequence similarities between *Klebsiella* phage Kpn_R1 (Seq1) and its closest relatives: *Klebsiella* phage vB_Kpn_IME260, *Klebsiella* phage Sugarland, and *Escherichia* phage N15. Conserved regions are depicted by connected syntenic blocks, with the strongest similarities shown in pink. The distinct structural organization of *Escherichia* phage N15 highlights its evolutionary divergence.

As shown in S4 Table, the whole-genome sequencing of the isolated phage demonstrated a 92.8% identity across 31.5% of the whole genome of its closest similar *Klebsiella* phage vB_Kpn_IME260. *Klebsiella* phage Sugarland (NC_042093) exhibited a mean identity of 92.3% across 30.1% of the genome length, indicating a modest degree of similarity with related phages. Sugarland is classified within the Pseudomonadota category, signifying a wider host range among Enterobacteriaceae members.

The comparison also recognized *Escherichia* phage N15 (NC_001901) as a similar phage, however it had a lower mean identity of 69.8% across merely 14% of the genome. Different from lytic phages such Sugarland, N15 is a completely defined temperate phage that may integrate into the bacterial genome as a linear plasmid. The somewhat low identity and genome coverage suggest that N15 is evolutionarily different even if it shares certain genetic elements with the isolated phage. These findings suggest that the isolated phage may adapt its genome for better survival or bacterial genome evolution via horizontal gene transfer technologies due to its evolutionary links and functional capacity. Comparative genomics will dominate future research to identify distinct genetic variables influencing host specificity and lytic activity.

### Klebsiella phage Kpn_R1 genome annotation

The 220 ORFs (S3 Table) in the annotated phage genome have been categorized into functional categories based on their anticipated functions. As shown in Table 7, hypothetical/uncharacterized proteins comprise the majority of the total ORFs, accounting for 48.34% (102). These proteins are not known to have any known functions or exhibit any significant similarities to known proteins. This implies that there are novel genes that are unique to phages and may play distinct functions in the phage lifecycle. Structural proteins constitute the second-largest group, which accounts for 18.01% (38) of the ORFs. This encompasses the primary capsid protein, tail fiber proteins, and portal proteins, all of which are crucial for the formation of virion structures. The significant presence of structural proteins highlights an accurate virion assembly for effective infection and replication. Proteins associated with DNA replication and metabolism comprise 11.85% (25) of the open reading frames (ORFs). DNA polymerase, helicase, exonuclease, and recombination mediators are all in this group. These components are essential for the preservation and replication of genomes. The phage’s ability to modify host DNA and facilitate effective genome replication is emphasized by the presence of these proteins. Transcriptional regulators, anti-termination proteins, and XRE family regulators comprise 2.84% (6) of the ORFs. It is probable that these proteins are involved in the regulation of gene expression throughout the phage lifecycle, which allows for the rapid activation or suppression of genes that are necessary for the various stages of infection.

**Table 7.**
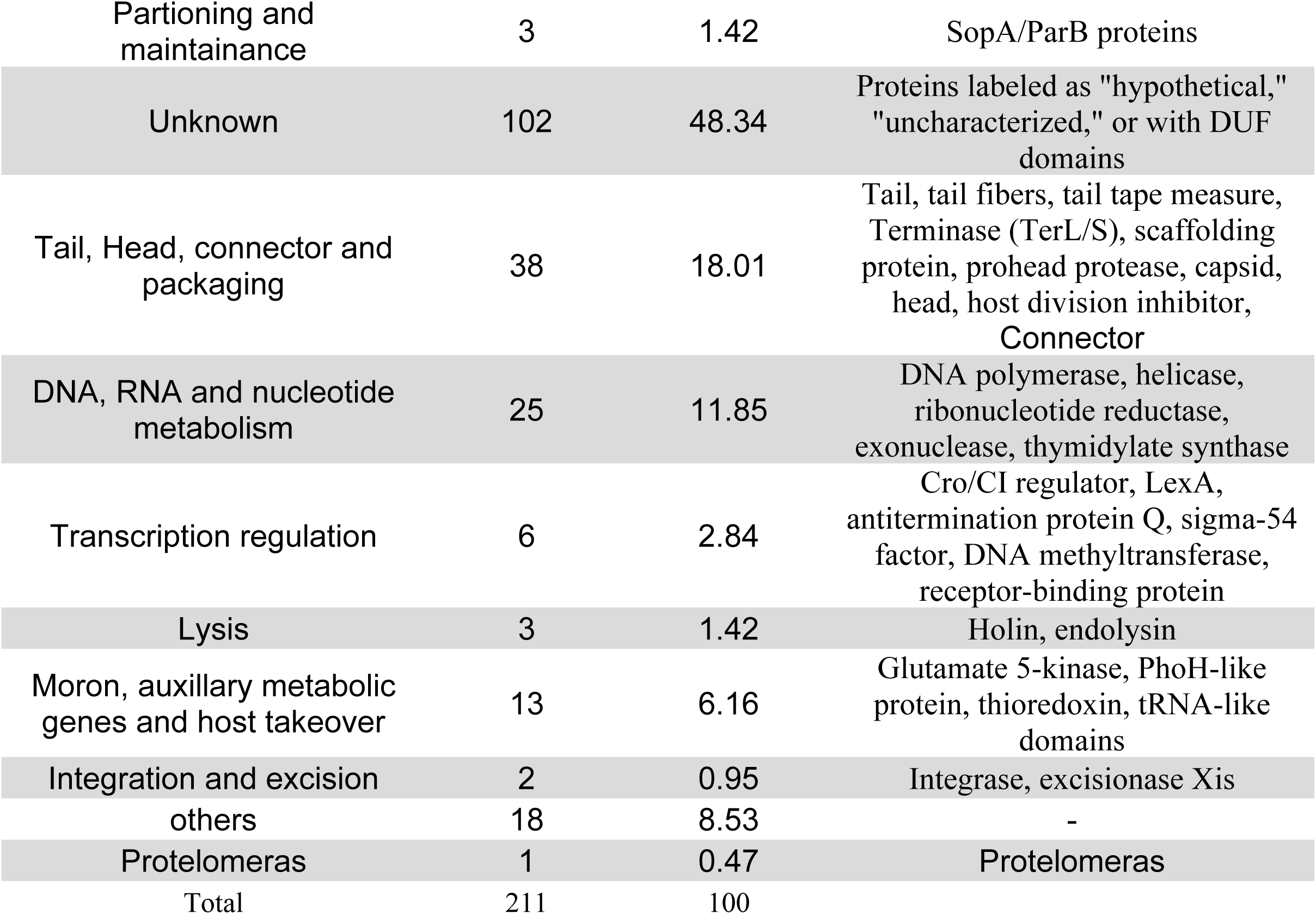
Functional Categorization of *Klebsiella* Phage Kpn_R1 Open Reading Frames (ORFs)

The host lysis group, which comprises endolysin, holin, and spanin proteins, comprises 1.42% (3) of the ORFs. These proteins are crucial for lysing the host cell to liberate freshly formed phage particles, signifying the concluding phase of the lytic cycle. The category of Moron, auxillary metabolic genes and host takeover comprising 6.16% (13) of the ORFs. Also included in the genome annotated ORFs, the ParA/B-like proteins comprising 1.42% (3). These proteins participate in plasmid partitioning with protelomerase presence, indicating a mechanism for the steady inheritance of the phage genome throughout cellular division, akin to the N15 phage and protelomerase reinforces this similarity, since protelomerase is characteristic of N15’s replication and maintenance approach, facilitating telomere resolution and linear genome preservation. The genome encodes a toxin-antitoxin (TA) system, especially a RelE/ParE family toxin (S3 Table). Plasmid maintenance, stress response, and persistence mechanisms are frequently implemented by TA systems. In phages, they may influence the behavior of host cells to promote phage survival or facilitate the stability of the prophage state. The phage genome exhibits a functional distribution that is typical of temperate phages, with a focus on host lysis, DNA replication, and structural assembly. The extensive unexplored biology is suggested by the significant predominance of putative proteins. However, the presence of partitioning proteins, protelomerase, and a toxin-antitoxin system suggests a reproductive and maintenance strategy similar to that of the N15 phage. In order to confirm the mechanisms of the phage’s lifecycle and to elucidate the functions of the potential proteins, additional experimental investigations are required.

## Discussion

The analysis of genome size, GC content, resistome, virulence determinants, gene transfere mechanisms and multilocus sequence typing (MLST) provides critical insights into the genetic diversity and evolutionary dynamics of bacterial pathogens, particularly for high-risk clones such as ST147. ST147, a globally disseminated high-risk clone of *K. pneumoniae,* has been associated with multidrug resistance and nosocomial infections, posing significant challenges to public health (44,45). In this study, we investigated the genomic characterization and MLST profile of ST147 isolate, to better understand its genetic characteristics In the current study, *K. pneumoniae* Kpn-R1 strain was isolated from an ICU patient’s sputum sample. The strain has a strong antibiotic resistance profile. The strain’s genome size was 5,835,643 bp, with a GC content of 56.82%, according to whole genome sequencing. The strain was identified as a high-risk ST 147 clone using Pasteur Institute MLST. The genome size of *K. pneumoniae* ST147 isolates is generally between 5.0 and 5.5 Mbp, consistent with prior findings (1). The average GC content of ST147 isolates in our investigation was 56.82%, which is within the recognized GC content range for *K. pneumoniae* (56-58%) (46). These genomic traits reflect the species’ flexibility and ability to accept foreign genetic material, such as plasmids and integrons, which typically carry antibiotic resistance genes (47). The genome’s stability and the persistence of resistance determinants in clinical settings may also be facilitated by the relatively high GC content.

The Gulf Cooperation Council (GCC) nations, in particular, have become hotspots for ST147 and its variations in the Middle East. ST147 is often isolated from patients in intensive care units (ICUs) in Saudi Arabia and has been linked to carbapenem-resistant *K. pneumoniae* (CRKP) outbreaks (48). ST147 has been found in clinical and environmental samples from the United Arab Emirates (UAE) and Oman, often in association with multidrug resistance (49,50). The high frequency of ST147 in the area may be due to a lack of infection control measures in specific healthcare institutions, excessive use of broad-spectrum antibiotics, and increased patient movement.

This research focuses on the XDR *K. pneumoniae* strain as the major danger it poses to public health calls for attention. Resistance profile of this strain is complex and influenced by 32 AMR genes spanning many classes of antibiotics. These genes encode resistance mechanisms showing the bacteria can avoid practically all medicines. These include Beta-Lactams, fluoroquinolones, aminoglycosides, tetracyclines, macrolides, sulfonamides, phenicols, polymyxins, and sulfonamides. According to Nordmann et al. (2011) and Logan and Weinstein (2017), there is a wide resistance pattern associated with the increasing incidence of *K. pneumoniae* bacteria that are resistant to several drugs. This trend is especially noticeable in healthcare institutions (2,51).

In the present work, SHV-11 beta-lactamase, NDM-1 carbapenemase, OmpK37, and MdtQ mediate beta-lactams resistance genes discovered in the genome of *K. pneumoniae.* SHV-11 beta-lactamase and NDM-1 carbapenemase are the primary enzymes that facilitate resistance to beta-lactams, including penicillins, cephalosporins, and carbapenems. These enzymes hydrolyze these antibiotics, thereby negating their efficacy (52). Furthermore, the migration of beta-lactams into the bacterial cell is restricted by mutations in porins (OmpK37, MdtQ), which reduce permeability (53). These pathways combined provide significant resistance to beta-lactams, including the last-resort carbapenems.

ST147 is a high-risk clone that has been extensively documented and is associated with carbapenem resistance, notably as a result of the production of carbapenemases like NDM-1 and OXA-48 (54). Studies undertaken in Saudi Arabia have highlighted the frequency of ST147 in clinical settings, with a significant percentage of isolates carrying the blaNDM-1 and blaOXA-48 genes (49,55). The spread of ST147 in this area is reason for worry since it highlights the importance of regional travel and healthcare networks in the spread of high-risk clones.

Resistance to fluoroquinolones, including ciprofloxacin (≥4 μg/mL) and levofloxacin (≥8 μg/mL), is facilitated by a synergy of efflux pumps (oqxA, acrA, marA) and mutations in the target enzymes DNA gyrase and topoisomerase IV (gyrA, parC) (56). These methods diminish intracellular drug concentrations and modify drug-binding sites, hence imparting significant resistance to this family of antibiotics.

The strain’s resistance to aminoglycosides, such as gentamicin (≥16 μg/mL) and amikacin (≥64 μg/mL), is mediated by 16S rRNA methyltransferase (armA) and aminoglycoside-modifying enzymes (aadA2, aac(6’)-Ib) (57). Aminoglycosides are rendered ineffective by these enzymes, which alter the antibiotic target or the substance itself. The strain’s capacity to resist this class of antibiotics is emphasized by the presence of numerous aminoglycoside resistance genes.

Efflux pumps (tet(A), tet(B)) and ribosomal protection proteins (tet(M)) mediate resistance to tetracyclines (tetracycline ≥16 μg/mL, tigecycline ≥2 μg/mL), which reduce intracellular drug concentrations or protect the ribosome from inhibition (58). Similarly, efflux transporters (msrE, mphE) and enzymatic inactivation (ereA) are mechanisms that have been increasingly observed in Gram-negative bacteria (59) that facilitate resistance to macrolides (erythromycin ≥8 μg/mL, azithromycin ≥16 μg/mL).

The strain’s resistance to trimethoprim (≥16 μg/mL) and sulfamethoxazole (≥256 μg/mL) is mediated by dihydropteroate synthase (sul1, sul2) and dihydrofolate reductase (dfrA12, dfrA14), respectively. The drug targets are altered by these enzymes, which prevent the inhibition of folate biosynthesis, a critical pathway for bacterial proliferation, thereby conferring resistance (60).

Efflux transporters (cmlA, floR) and enzymatic inactivation (catA1) are responsible for chloramphenicol resistance (≥32 μg/mL) (61). Similarly, fosfomycin thiol transferase (fosA5) is responsible for the inactivation of the antibiotic, resulting in resistance to fosfomycin (≥64 μg/mL) (62). The strain’s capacity to evade antibiotics that target a variety of cellular processes is underscored by these mechanisms.

Phosphoethanolamine transferase (arnT, eptB) and mutations in the mgrB gene are associated with resistance to colistin and polymyxin B (≥4 μg/mL). These mutations modify the lipid A component of lipopolysaccharide, thereby reducing the binding affinity of polymyxins (63). This mechanism is particularly alarming, as polymyxins are frequently employed as last-resort antibiotics to treat XDR Gram-negative infections.

The efflux transporters (qacEdelta1, leuO) that are frequently associated with biocide resistance in Gram-negative bacteria facilitate the strain’s resistance to quaternary ammonium compounds (≥50 μg/mL) (64). The strain’s persistence in hospital environments, where biocides are frequently employed for disinfection, may be influenced by this resistance mechanism.

In addition to its extensive resistance profile, the strain possessed a number of virulence factors that exacerbated its pathogenesis. The E. coli common pilus (ECP) gene cluster’s presence (98.31–100% coverage, 85.28–90.07% identity) implicates the strain’s capacity to adhere to host tissues and form biofilms, which is a critical factor in the establishment of infections (65). Enterobactin genes (88.72–99.33% coverage, 80.00–82.65% identity) and the yersiniabactin biosynthetic gene cluster (100% coverage, 97.62–99.95% identity) suggest that the strain is capable of acquiring iron, a critical nutrient for bacterial survival and virulence in the host (66). Outer membrane protein A has been demonstrated to safeguard microorganisms from host immune responses (67), which further supports the notion that the ompA gene has immune evasion capabilities (100% coverage, 83.75% identity). These virulence factors collectively contribute to the strain’s capacity to cause severe infections, particularly in immunocompromised patients.

According to Kaptive analysis, the K64 and O2a serotypes of the strain are in accordance with prior reports of XDR *K. pneumoniae* isolates that are linked to healthcare-associated infections (68). Potential genetic exchange and antigenic diversity are suggested by the presence of additional genes associated with other serotypes, including KL40, KL150, and KL4. This may facilitate persistence and immune evasion within the host. The strain’s ability to promote horizontal gene transfer (HGT) and the spread of antibiotic resistance genes is highlighted by its plasmid content, which includes IncHI1B_1_pNDM-MAR, IncFIB (K)_1_Kpn3, and IncFIB (pKPHS1)_1_pKPHS1 replicons (69). The strain’s capacity to produce bacteriocins is indicated by the presence of colicin genes (Col440I_1, Col (BS512)_1, ColpVC_1), which may enhance its survival in polymicrobial environments and inhibit the growth of competing bacterial strains (70).

The strain exhibits the Type I-E CRISPR-Cas gene cluster, conferring adaptive immunity against foreign genetic elements, including plasmids and phages (71). This system may influence the strain’s capacity to acquire and retain resistance genes by regulating its interactions with phages and other mobile genetic elements. Eleven prophage regions are present which are often harbor genes that enhance bacterial virulence and antibiotic resistance, highlighting the genetic diversity of strains and their potential for horizontal gene transfer (5).

The annotated phage genome analyzed in this study reveals a complex and multifaceted genetic architecture, characteristic of temperate phages. The presence of a linear genome, protelomerase, and ParA/B-like partitioning proteins strongly suggests a replication and maintenance strategy similar to that of the N15 phage, a well-studied temperate phage with a linear plasmid prophage lifestyle (7). The N15 phage is known for its unique ability to maintain its genome as a linear plasmid in the lysogenic state, facilitated by the action of protelomerase (TelN) and partitioning proteins (9). The protelomerase identified in this study shares 93.3% identity with known protelomerases, further supporting this resemblance. This finding aligns with previous studies that highlight the role of protelomerase in resolving telomeres and maintaining linear genomes in phage systems (7,72).

The toxin-antitoxin (TA) system identified in the genome belongs to the RelE/ParE family, which is often associated with plasmid maintenance, stress response, and persistence mechanisms (73). In phage genomes, TA systems are thought to stabilize the prophage state and modulate host cell behavior to favor phage survival (74). The presence of a TA system in this phage suggests a potential role in lysogeny maintenance or stress adaptation, which could be further explored through experimental studies.

The UmuD-like protein identified in the genome is part of the SOS response system, which is involved in DNA damage repair and mutagenesis (75). In phages, UmuD-like proteins may facilitate genome replication under stress conditions, such as exposure to UV radiation or chemical mutagens. This finding is consistent with studies on other phages, where UmuD homologs have been shown to enhance survival in hostile environments (76).

The structural proteins identified in the genome, including major capsid proteins, tail fibers, and portal proteins, are essential for virion assembly and host recognition. These proteins are highly conserved among phages and play a critical role in the infection cycle (77). The high proportion of structural proteins (11.8%) in this genome underscores their importance in the phage lifecycle, particularly in the assembly of infectious particles.

The hypothetical proteins constitute the largest category (67.5%) of the ORFs, highlighting the significant proportion of uncharacterized genes in the phage genome. This is not uncommon in phage genomics, as many phage genes lack homology to known proteins and may encode novel functions (78). The high number of hypothetical proteins suggests that this phage may possess unique adaptations or mechanisms that have yet to be discovered. Future functional studies, such as transcriptomics and proteomics, could shed light on the roles of these proteins.

The regulatory proteins identified in the genome, including transcriptional regulators and anti-termination proteins, are likely involved in controlling gene expression during the phage lifecycle. These proteins ensure the timely activation or repression of genes required for different stages of infection, such as lysogeny and lytic replication (79). The presence of XRE family transcriptional regulators further supports the phage’s ability to modulate host gene expression, a common strategy among temperate phages (80).

The phage genome analyzed in this study exhibits a functional distribution typical of temperate phages, with a strong emphasis on structural assembly, DNA replication, and host lysis. The presence of partitioning proteins, protelomerase, and a toxin-antitoxin system suggests a replication and maintenance strategy similar to that of the N15 phage. However, the high proportion of hypothetical proteins indicates significant unexplored biology, highlighting the need for further experimental studies to elucidate the roles of these genes and confirm the phage’s lifecycle mechanisms.

## Conclusions

The XDR *K. pneumoniae* Kpn_R1 strain analyzed in this study is a significant pathogen due to its extensive array of virulence factors and resistance characteristics, which are governed by 32 AMR genes. The strain’s ability to evade the human immune system and antibiotics underscores the urgent requirement for alternative therapeutic strategies, such as phage therapy or combination treatments, to combat infections caused by these strains. Future research should focus on elucidating the functional roles of the strain’s virulence factors and resistance mechanisms, alongside exploring the potential of CRISPR-Cas systems and prophages as targets for novel antimicrobial treatments.

The linear genome structure, protelomerase-mediated replication, and partitioning mechanism of the annotated phage closely resemble those of the N15 phage. This may indicate its affiliation with the N15-like phage group, as these characteristics suggest a replication and maintenance strategy akin to that of N15. Additional research will elucidate the biology of this phage and its relationship to N15, encompassing functional characterization of the putative proteins and experimental validation of the replication process.

## Material and methods

### Culture Conditions and Bacterial Strains

Intensive care units (ICUs) extensively drug resistant (XDR) clinical strain of *K. pneumoniae* was isolated from sputum sample. Bacterial identification and the antibiotic susceptibility profiles and MICs of the isolates were assessed by VITEK 2 system (VITEK® 2, BIOMÉRIEUX, France). Bacterial strains were cultivated in Luria-Bertani (LB) broth or on LB agar plates at 37°C and preserved at −80°C in glycerol stocks for long-term storage.

### Bacteriophage Isolation and Purification

Environmental samples (hospital effluents) were collected in a sterile 1 L sterile glass bottle for phage isolation. The enrichment method was used, where 10 mL of sample was mixed with 10 mL of LB broth containing a bacterial host, incubated at 37°C for 24 hours with shaking, and filtered through a 0.22 µm membrane (14). The resulting phage suspension was serially diluted and plated on soft LB agar containing the host bacteria to detect plaque formation. Single plaques were picked, purified by repeated plaque assays, and stored at 4°C.

### Phage Characterization

Phage morphology was observed using Transmission Electron Microscopy (TEM) according to standard procedures (15). Briefly, a drop of the purified phage suspension was placed on a carbon-coated copper grid, stained with 2% uranyl acetate, and examined using a JEOL JEM-1400 electron microscope at 120 kV.

The host range of the isolated phage was determined by performing plaque assays against a panel of clinical *K. pneumoniae* strains. The strains were cultured overnight and mixed with soft agar containing the host bacterial strains, and plaque formation was observed to determine the host range of the phage (16).

### DNA isolation and Whole-Genome Sequencing (WGS)

The genomic DNA of the isolated phage was extracted using the Phage DNA Isolation Kit (Norgen Biotek) according to the manufacturer’s protocol. Whereas the bacterial DNA was isolated using Qiagen blood and tissue DNA extraction kit. The DNA concentration was measured using a Nanodrop spectrophotometer (Thermo Fisher Scientific, USA).

Macrogen Korea performed whole-genome sequencing (WGS) using the Illumina NovaSeq 6000 system. Following manufacturer directions, genomic DNA libraries were built using the TruSeq Nano DNA (350) kit. A library size check was part of the preparation for the library; the Agilent Technologies 2100 Bioanalyzer with a DNA 1000 chip validated the size distribution of PCR-enriched fragments. On the Illumina sequencing technology, optimum cluster densities were obtained by means of DNA library template quantification. Libraries were measured using qPCR applied under the Illumina qPCR Quantification Protocol. Libraries’ concentration and size distribution were confirmed using the TapeStation D1000 Screen Tape. Using the Illumina NovaSeq 6000 platform, the libraries were sequenced producing excellent paired-end reads. The phage and bacterial genome were built from processed and examined sequencing data. Raw reads were pre-processed using Trimmomatic for quality trimming (17). The cleaned reads were then assembled into contigs using SPAdes (18) with default settings.

### Phage WGS bioinformatics analysis

Raw sequencing data were assembled using SPAdes (v3.15.5) (18) then FastaQC was used to evaluate quality. Genome completeness was ascertained using CheckV (v1.0.1) (19), and structural features were projected using PhageTerm (v4.0.0) (20). Functional annotations included phage lifestyle prediction using Bacphlip (v0.9.6) (21) and protein classification using PhANNs (22). Comparative genomics was run using Pyani (v0.2.12) (23) and Clinker (v0.0.24) (24). AMRFinderPlus (v3.12.8) (25) and VirulenceFinder (v2.0.4) (26) separately verified antimicrobial resistance and virulence genes. MASH distances were used to show the genome in CGView (v1.1.1) (27) against the Millardlab Phage Database (28). Phylogenetic trees was generated using Viptree (29). Every tool is open source, hence once released data will be available in public repositories.

### Bacterial WGS bioinformatics analysis

Raw data from whole-genome sequencing (WGS) were integrated using Proksee Assemble (v1.3.0) (30). The assembled contigs were annotated using Prokka (v1.2.0), a rapid tool for prokaryotic genome annotation that detects open reading frames (ORFs), tRNA, rRNA, and other genomic features (31). Open reading frames (ORFs) were predicted using the ORFs (v1.0.1) program, which is designed to locate potential protein-coding regions in the genome. Map Builder (v2.0.5) generated a CGView JSON file for showing the completed genome from GenBank formats. To rank BLAST tracks based on similarity and color BLAST features based on percent identity, the BLAST Formatter (v1.0.3) was used, assisting in the identification of conserved and divergent regions.

Alien Hunter (v1.1.0) was used to forecast likely horizontal gene transfer (HGT) events based on atypical nucleotide compositions that indicate alien DNA acquisition (32). To compare the genome with closely related species and determine taxonomic connections, the whole-genome average nucleotide identity (ANI) was calculated using FastANI (v1.1.0) (33). The Comprehensive Antibiotic Resistance Database (CARD) Resistance Gene Identifier (RGI) (v1.2.1) was used to identify antibiotic resistance genes. Based on the CARD database, this tool anticipates resistance genes and mutations, giving information on the genome’s antibiotic resistance profile (34).

VirSorter (v1.1.1) was used to find viral genomes (35). Phigaro (v1.0.1) has been annotated in prophage regions (36). Mobile OG-db (v1.1.3), a database and tool designed for the identification of Mobile Genetic Elements (MGEs), including plasmids, transposons, and integrons (37). Using the CRISpen/Cas Finder (v1.1.0) tool, was used to locate CRISPER arrays and associated Cas proteins, offering insights into the genome’s adaptive immune system (38). The Proksee tool displayed and organized all genetic traits. All visible tracks on the genome map were documented in a detailed figure caption created using the Track List Caption (v1.2.0) (30).

Using Kaptive Web (39), capsule (K) and lipopolysaccharide (O) serotypes in *K. pneumoniae* genomes were predicted. Sequence-based typing and known *Klebsiella* strain comparisons using the PubMLST Klebsiella database (40). Virulence factors are identified using the 2019 Virulence Factor Database (VFDB) (41). Barrnap helped to pull out the 16S rRNA gene sequence (42). By use of NCBI BLASTn database comparison, closely similar species were identified. A phylogenetic tree based on the 16S rRNA sequence and other conserved markers was built using MEGA X (43), therefore exposing the evolutionary links of the isolate.

### Statistical analysis

Statistical analyses were conducted to evaluate the significance of observed patterns in the genomic and phenotypic data. Descriptive statistics were used to summarize key metrics, such as genome size, GC content, and gene coverage. Comparative analyses were performed to assess genetic similarity and divergence, with significance determined using p-values and distance metrics. Hypothesis testing was applied to evaluate associations between genetic features (e.g., resistance genes, virulence factors) and phenotypic outcomes (e.g., antibiotic resistance profiles). Statistical significance was defined at a threshold of p < 0.05. All analyses were performed using R (R Core Team, 2023) and Python (Python Software Foundation, 2023), with appropriate statistical packages for data visualization, hypothesis testing, and correlation analysis.

## Acknowledgments

I thank King Abdullah International Medical Research Center (KAIMRC) in Riyadh, Saudi Arabia for fund this project.

## Ethical research consideration

This study was conducted in conformity with ethical policies concerning the use of microbial genetic data. Examined *K. pneumoniae* strain came from a clinical sample taken under informed permission and approved by the relevant institutional review board (IRB) NRC21R/053/02. Patients’ right to privacy and confidentiality was fiercely maintained to guarantee that the research included no personally identifying data whatsoever. Complying with relevant national and international ethical research norms, the study adhered to the criteria established in the Declaration of Helsinki.

